# *In-silico* study of fatty acid biosynthesis pathway enzymes in microalga *Scenedesmus*

**DOI:** 10.1101/2024.05.01.592012

**Authors:** Harshit Kumar Sharma, Ma. Belén Velázquez, Noelia Marchetti, Ma. Victoria Busi, Julieta Barchiesi, Chitralekha Nag Dasgupta

## Abstract

*Scenedesmus quadricauda* is an important, rapidly growing, freshwater microalga explored as a source of alternative fuel because of its significantly high lipid content. However, the molecular basis of fatty acid biosynthesis is scarcely elucidated not only in *S. quadricauda* but also in *Scenedesmus* as a whole. This study aims to understand the 3D model structure of enzymes involved in fatty acid synthesis, and their catalytic sites compared to other *Scenedesmus* species. The first genome sequence *S. quadricauda* was carried initially in our isolate LWG002611 (GenBank ID NNCB00000000). However, till date, no study has been carried out on the 3D modelling and identifying the catalytic sites of fatty acid biosynthesis pathway enzymes. Mining the genome sequence of our isolated *S. quadricauda* LWG002611 as well as other *Scenedesmus* sequences taken from PhycoCosm and NCBI, we identified sequences of the crucial enzymes involved in fatty acid biosynthesis pathways such as acetyl-CoA carboxylase (ACC), malonyl-CoA:ACP transacylase (MAT) and fatty-acyl thioesterases (FAT) for comparative study on homology, catalytic sites, domains, and protein 3D models. Detailed comparative analyses of these identified enzymes were carried out using various bioinformatics tools. Which demonstrated highly significant sequence similarity with homologs of bacteria as well as with homologs of lower groups of eukaryotes suggesting an evolutionary linkage with them. The molecular modeling and 3D structures of the chloroplastic enzymes by AlphaFold Multimer revealed the overall structural orientation and well-conserved catalytic residues. On the other hand, biotin protein ligase and cytosolic acetyl-CoA carboxylase isoforms presented some significant differences with respect to the previously reported protein models. The conserved domain suggests the preservation of the fatty acid biosynthesis pathway in *Scenedesmaceae* family. However, some contrasting results, unique sequences and binding sites of some enzymes in *S. quadricauda* may have a significant role in higher lipid accumulation than the other species. Our analysis describes some specific features in *S. quadricauda* fatty acid synthesis enzymes that could open up the scope of further analysis of these enzymes.

## 1. Introduction

Climate change and scarcity of fossil fuels triggered researchers to identify biological solutions as an alternative to fuel and other bioproducts. Compared to other biofuel resources, algal biofuel is more attractive in many aspects, but a critical hurdle is to achieve the desired quantities of biomass and lipid concurrently^1,2^. Most of the studies used nutrient stress to enhance increased lipid production but ended up reducing the growth rate which affects overall biomass and lipid productivity^3^ as the correlation between lipid biosynthesis and microalgal growth has not been fully elucidated. Understanding the detailed mechanism of lipid biosynthesis in microalgae may provide a clue to manipulate the pathway for improving lipid yield without compromising on growth characteristics.

A key enzyme of fatty acids (FAs) biosynthesis is acetyl-CoA carboxylase (ACC) present in two distinct forms in algae; the homomeric form present in the cytosol and heteromeric form which is present majorly in plastid^4^. The first committed step for *de novo* FAs synthesis is the ATP-dependent carboxylation of acetyl-CoA to produce malonyl-CoA catalyzed by chloroplastic heteromeric ACC within the stroma^4^. The heteromeric ACC comprises four subunits named biotin carboxylase (BC), biotin carboxyl carrier protein (BCCP), and α- and β-carboxyltransferases (α- and β-CT)^5^. The Cytosolic ACC is known to provide malonyl-CoA for the elongation of FAs in the endoplasmic reticulum (ER)^6,7^. *In-silico* study of *Chlamydomonas reinhardtii* genome, revealed the occurrence of both types of ACC but the cellular location has not been verified^8^. The malonyl-CoA generated by chloroplastic ACC in the stroma enters the *de novo* FAs synthesis pathway and is attached to acyl carrier protein (ACP) to produce malonyl-acyl carrier protein by the enzyme malonyl-CoA:ACP transacylase (MAT)^5^, which is the starting point of multienzyme based elongation of acyl groups^5^. Chain elongation terminates by fatty-acyl thioesterases (FAT) which hydrolyses acyl-ACP to form non-esterified FAs and ACP. Based on sequences and substrate specificities, two types of FATs have been identified, FatA and FatB, which determine the types and quantities of FAs^8^. The preferred substrate of FatA is 18:1-ACP, whereas FatB primarily hydrolyzes the saturated 8-18 carbons-ACP^8^. In FA biosynthesis pathway ACC, MAT, and FAT are promising targets for genetic manipulation. In-depth bioinformatic analysis may provide necessary information to tailor algae for higher FAs production. The first *in-silico* study on FAs biosynthesis enzymes was carried out in model green alga *C. reinhardtii*^6,9^ and with the advancement of genome sequencing, many algal genomes have been sequenced and subjected to mining of genome for biochemical pathways related to lipid metabolism^10–12^. Recently, microalgae *Scenedesmus* has gained a lot of attention due to its biofuel production potential^10,13^. Some substantial work has been carried out to sequence its genome and transcriptomes; furthermore, the triacylglycerol (TAG) biosynthesis pathway has been reconstructed ^10,14–18^. Despite numerous studies on the lipid biosynthesis pathway of *Scenedesmus*, there remains a notable absence of information regarding protein structures, conserved motifs, catalytic sites, and the mechanisms of action for many of the crucial enzymes. Particularly there is no bioinformatic analysis has been carried out on *S. quadricauda,* which is a very important microalga in the field of biofuel production.

In the present study, we have performed a comparative bioinformatic analysis of ACC, MAT, and FAT sequences identified from our previous published genome sequence of *S. quadricauda* LWG002611 (NCBI accession no. NNCB00000000) a highly oleaginous microalga^10^ with other species of *Scenedesmus* such as *S. obliquus* UTEX 3031, *S. obliquus* UTEX 393, *S. obliquus* UTEX 393 v 2.0, *S. obliquus* var. UTEX 1450 v1.0, *S. obliquus* var. UTEX2630 v1.0, *S. obliquus* var. DOE0013 v1.0, *Scenedesmus* sp. NREL 46B-D3, *Scenedesmus* sp. NREL 46B-D3 v1.0 and *Scenedesmus* sp. PABB004 taken from PhycoCosm and NCBI GenBank portal. Homology study of conserved domains of ACC, FAT, and MAT of *S. quadricauda* has been carried out for first time along with the three-dimensional (3D) modeling of enzymes. We have compared and demonstrated the binding sites and catalytic residues of fatty acid biosynthesis pathway enzymes in *S. quadricauda* LWG002611 and *S. obliquus* UTEX 3031.

## 2 Methods

### 2.1 Sequence search, alignment and homology analysis

The enzymes of TAG biosynthesis pathway acetyl-CoA carboxylase (ACC), malonyl-CoA:ACP transacylase (MAT), fatty acyl-ACP thioesterase (FAT) of *S. quadricauda* LWG002611 were taken from our previous results (NCBI accession no. NNCB00000000)^10^ and various species of family *Scenedesmaceae* were identified and taken from PhycoCosm^12^ and NCBI (https://www.ncbi.nlm.nih.gov/): *S. obliquus* UTEX 3031, *S. obliquus* UTEX 393, *S. obliquus* UTEX 393 v 2.0, *S. obliquus* var. UTEX 1450 v1.0, *S. obliquus* var. UTEX2630 v1.0, *S. obliquus* var. DOE0013 v1.0, *Scenedesmus* sp. NREL 46B-D3, *Scenedesmus* sp. NREL 46B-D3 v1.0 and *Scenedesmus* sp. PABB004. Their gene ID/protein ID/accession numbers are shown in Table 1. The sequences were checked for similarity using BlastN and BlastP^19^. The conserved domain were identified by InterPro (https://www.ebi.ac.uk/interpro/)^20^ and transit peptides (TP) for cell location were identified by TargetP 2.0^21^ (https://services.healthtech.dtu.dk/services/TargetP-2.0/), PredAlgo^22^ and SignalP 5.0 (https://services.healthtech.dtu.dk/services/SignalP-5.0/). Protein folding homology of TAG biosynthesis pathway enzymes with crystal structures of Protein Data Bank (RCSB) enzymes were analyzed by Phyre2 (http://www.sbg.bio.ic.ac.uk/~phyre2/html/page.cgi?id=index) or orthoDB (https://www.orthodb.org/?gene=A0A0D2K6G4). The amino acid sequences were aligned by Clustal Omega^23^ and visualized with ESPript 3.0^24^

**Table 1:**
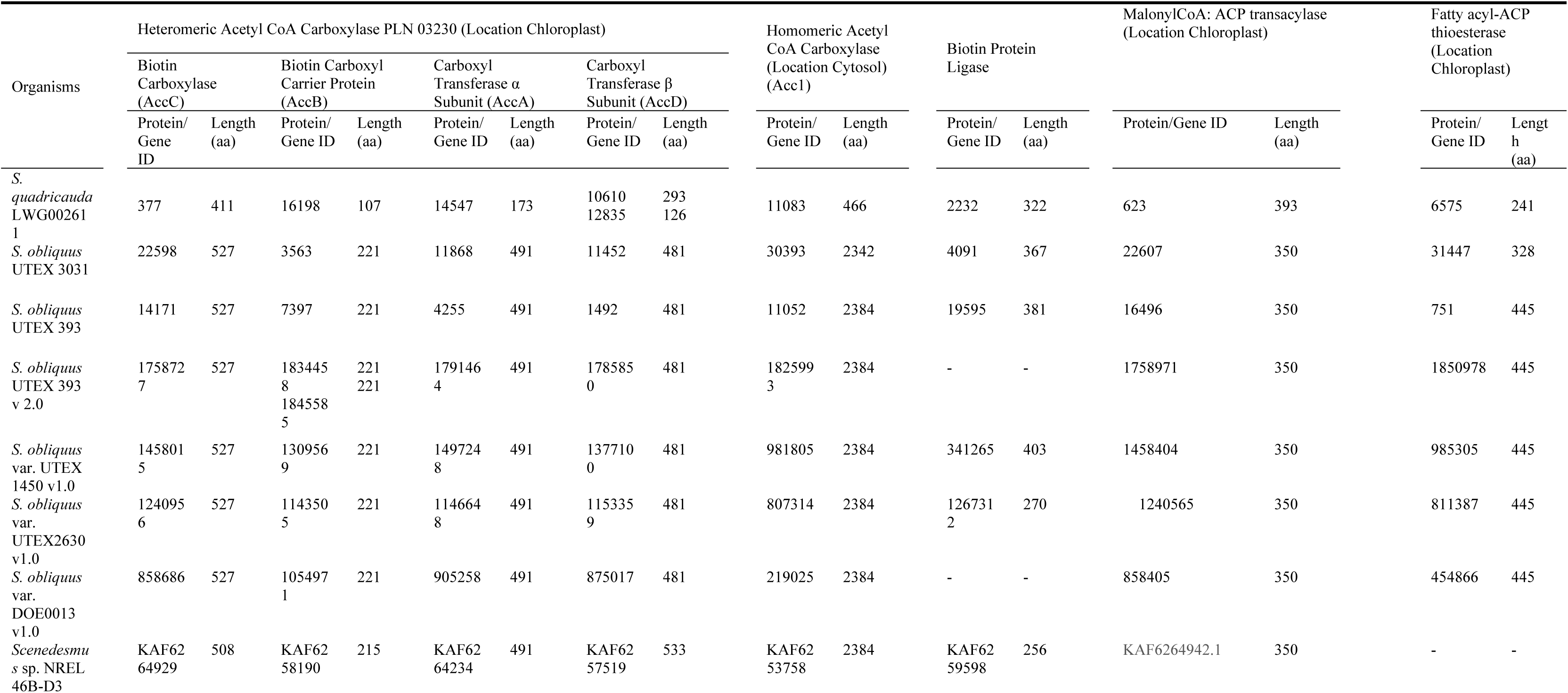
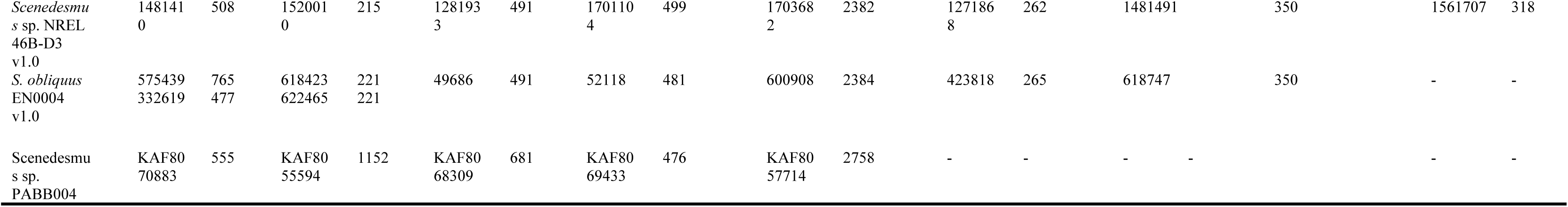
Fatty acid biosynthesis pathway enzymes (identified and taken from draft genome of S. quadricauda LWG002611, PhycoCosm and NCBI)

### 2.2 Protein 3D modeling

The 3D structure prediction of the studied proteins was generated in ColabFold using its interface. The MMseqs2 method was selected in ColabFold to generate the MSAs, the amber relaxation of the model was disabled and the unpaired MSA prediction was generated^25^. Protein complex prediction was done with AlphaFold-Multimer using the Cosmic platform, AlphaFold Multimer is an extension of AlphaFold2 that has been specifically built to predict protein-protein complexes with high accuracy^26^. The structures were visualized using the PyMOL Molecular Graphics System, Version 1.2r3pre, Schrödinger, LLC. Homology modeling was accomplished with the SWISS_MODEL server (https://swissmodel.expasy.org/) and default settings as an alternative method for short sequences.

## 3 Results and Discussion

### 3.1 *In-silico* analysis of Acetyl-CoA carboxylase (ACC)

We have performed an analysis of the fatty acid biosynthesis pathway in the draft genome of *S. quadricauda* LWG002611 (GenBank ID NNCB00000000) sequenced by us in our previous study^10^ and identified heteromeric ACC components BC (Sq377), BCCP (Sq16198), α-CT (Sq1497248), β-CT (Sq10610, Sq12835) and homomeric ACC (Sq11083) in the genome sequence^10^ (Table 1). A comparative study of *S. quadricauda* LWG002611 sequences with other *Scenedesmus* sequences tabulated in Table 1 was performed using different bioinformatic tools.

Analysis of sequences by Phyre2 and SWISS MODEL and alignments with Protein Data Bank (PDB) molecules revealed the homology of highly conserved motifs and conserved catalytic sites of the enzymes (Table 1, 2, Fig. 1).

**Fig 1:**
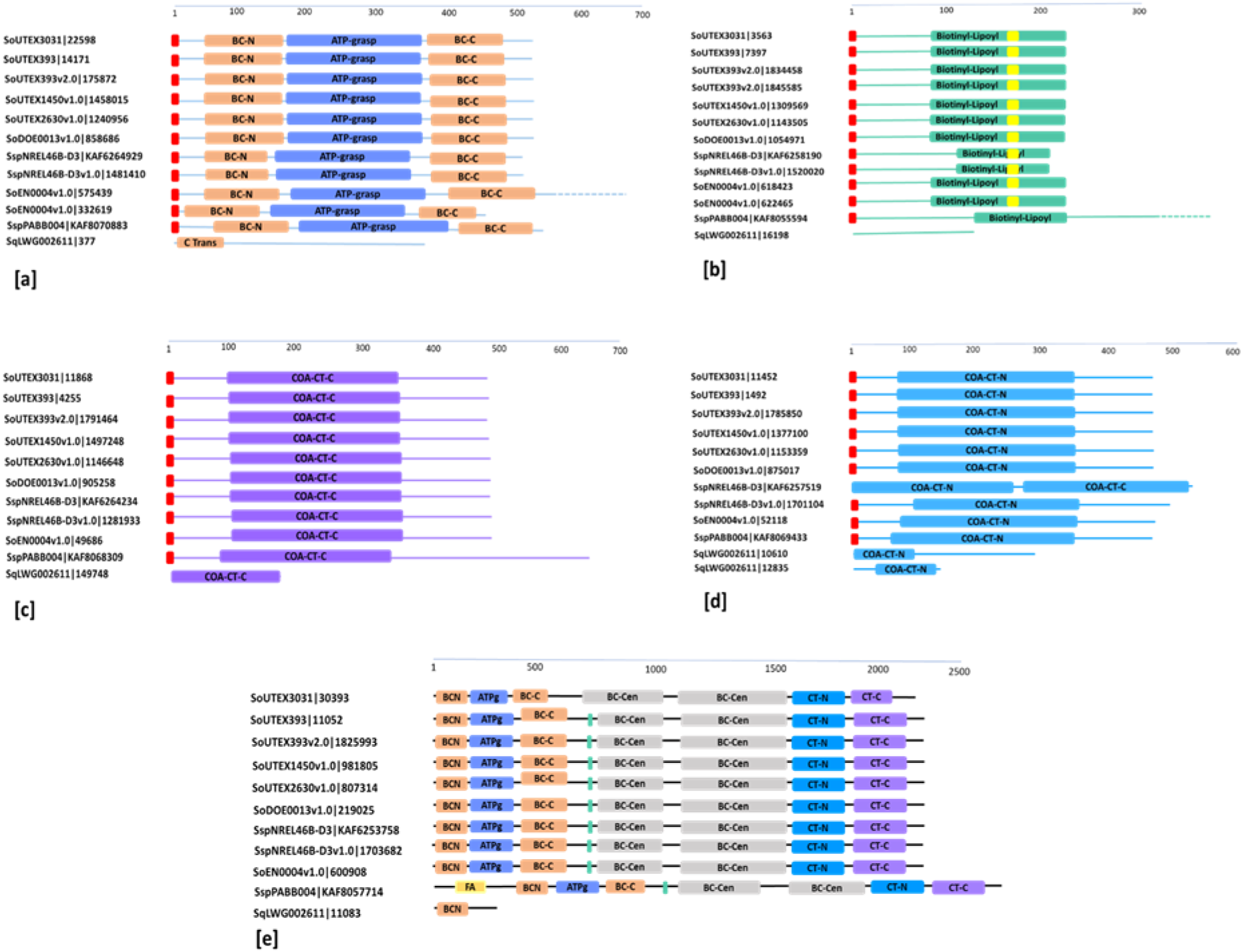
Protein domain architecture of ACC showing all InterPro predicted domains, red squares represent signal peptides. **(a)** N- and C-terminal domains of BCs are represented by orange rectangles and ATP grasp domains are represented by blue rectangles; **(b)** Biotinyl/Lipoyl domain of BCCPs are represented by green rectangular bar and biotin-binding sites are represented by yellow squares; **(c)** C-terminal domain of α-CTs are represented by purple rectangular bar **(d)** N-terminal domain of β-CTs are represented by light blue rectangular bar; **(e)** BC, ATP grapes, CT domains of homomeric ACCs on a single peptide are represented by corresponding colours.

*S. quadricauda* BC (Sq377) sequence showed homology with eukaryotic *Saccharomyces cerevisiae* (yeast) ACC however other *Scenedesmus* BC sequences showed homology with *E. coli* or *S. aureus* proteins. In domain study we have found only the carboxyl transferase domain in Sq377 whereas others BC proteins showed N- and C-terminal domain and ATP grasp domain (Fig. 1a). When we search for a template on PDB we find homology with the structure of thermophilic filamentous fungus *C. thermophilum* 5I6I (29.38% identity, covering 79%) (Fig. 3d). When the BC Sq377 3D SWISS-MODEL homodimer (GMQE of 0.51%) was superposed onto the 5I6I BC PDB structure we have found superposition only with the carboxyl transferase domain (Fig 3d). In this structure superpositions, the C-terminal region is defined as the C-terminal CoA carboxyltransferase and coincides with the model, spanning from residue 1486 to residue 2196.

**Fig 2:**
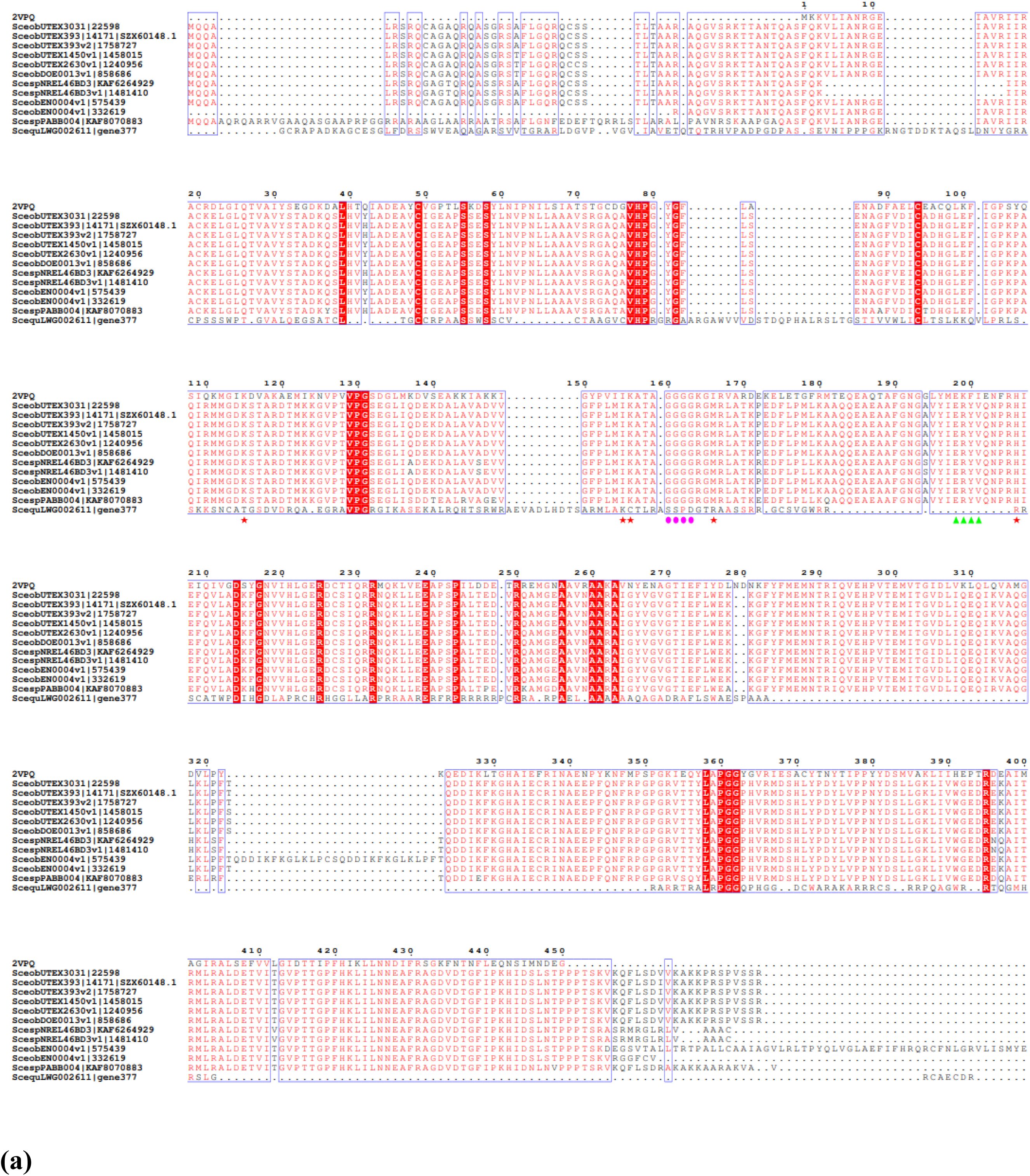
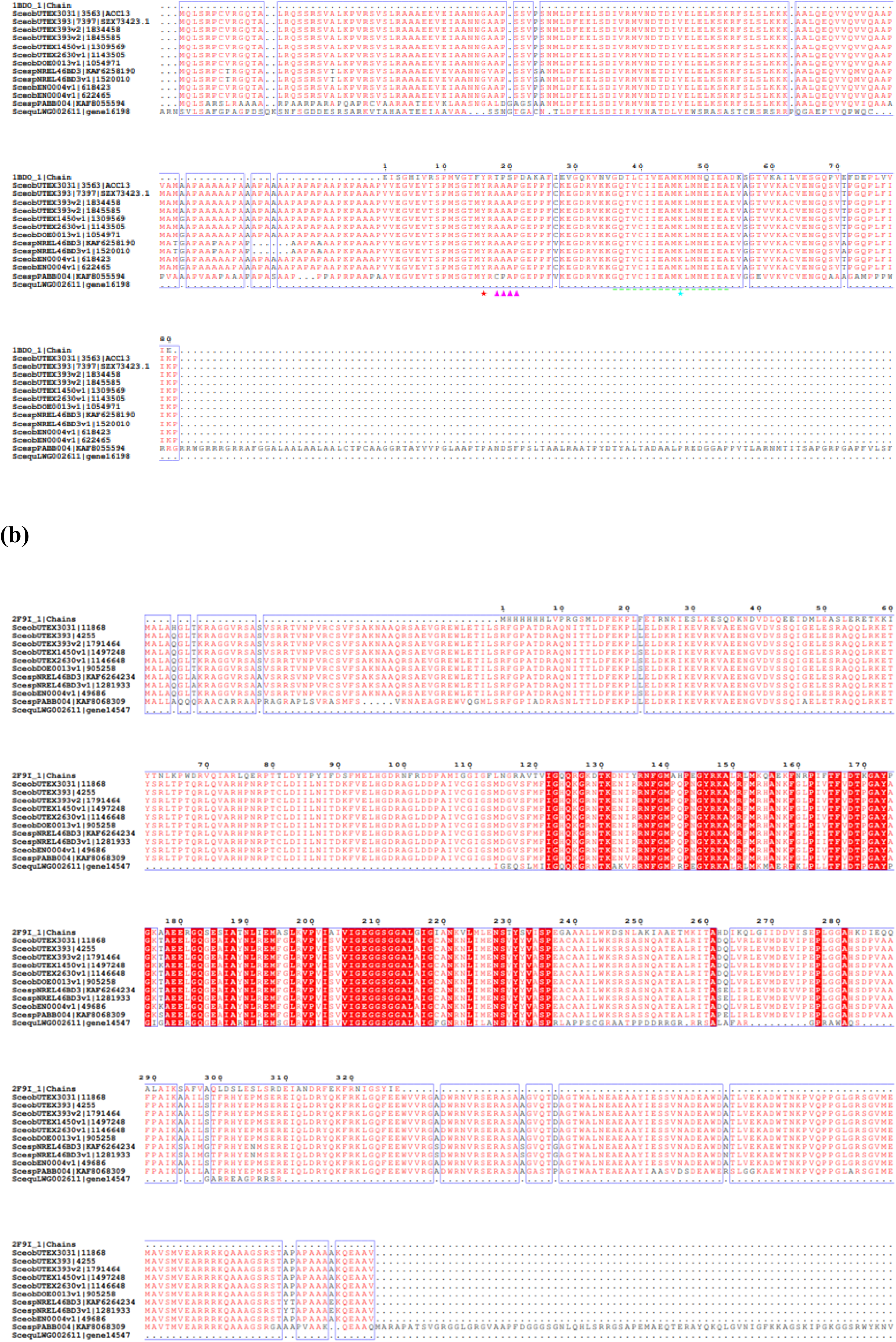
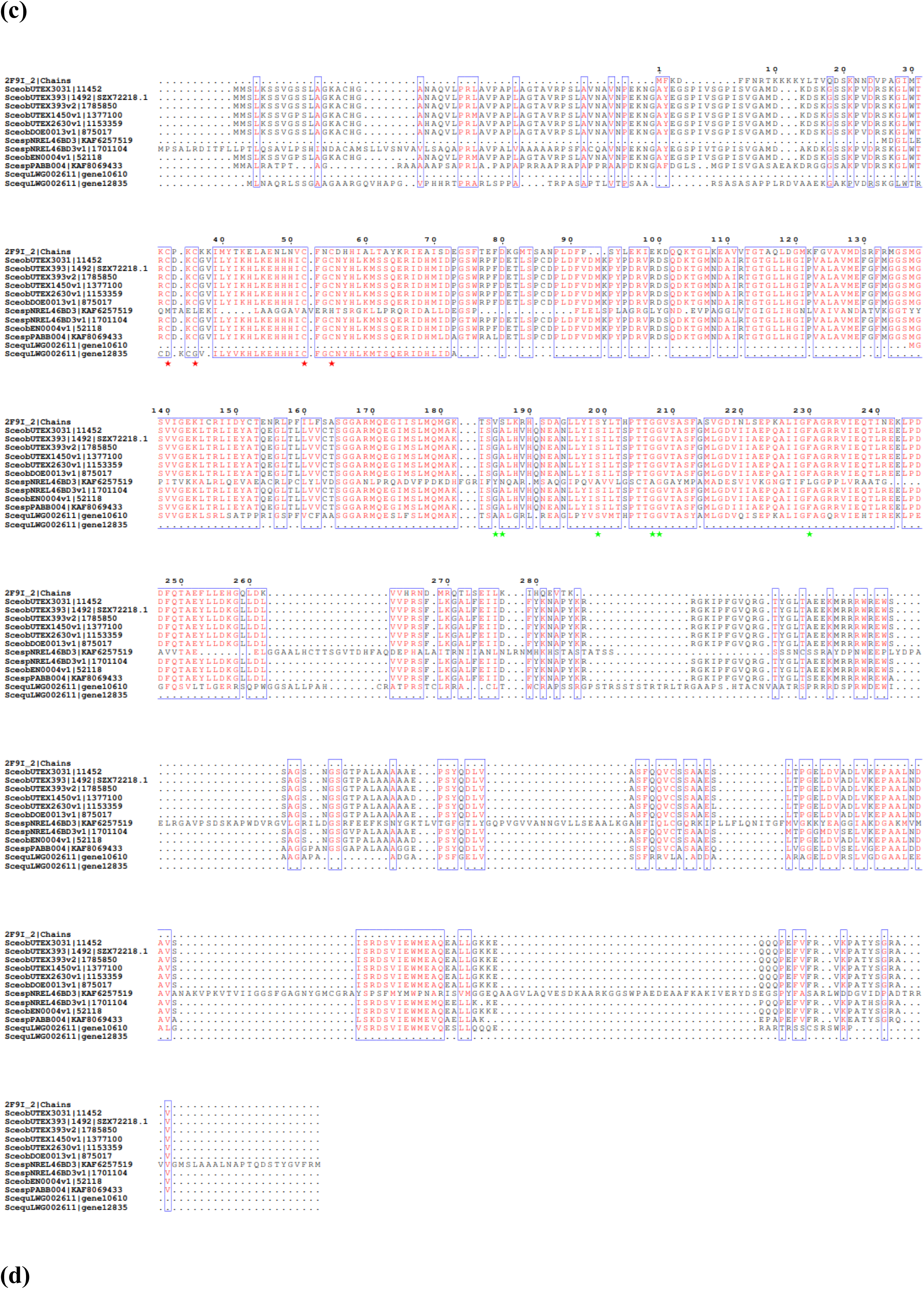
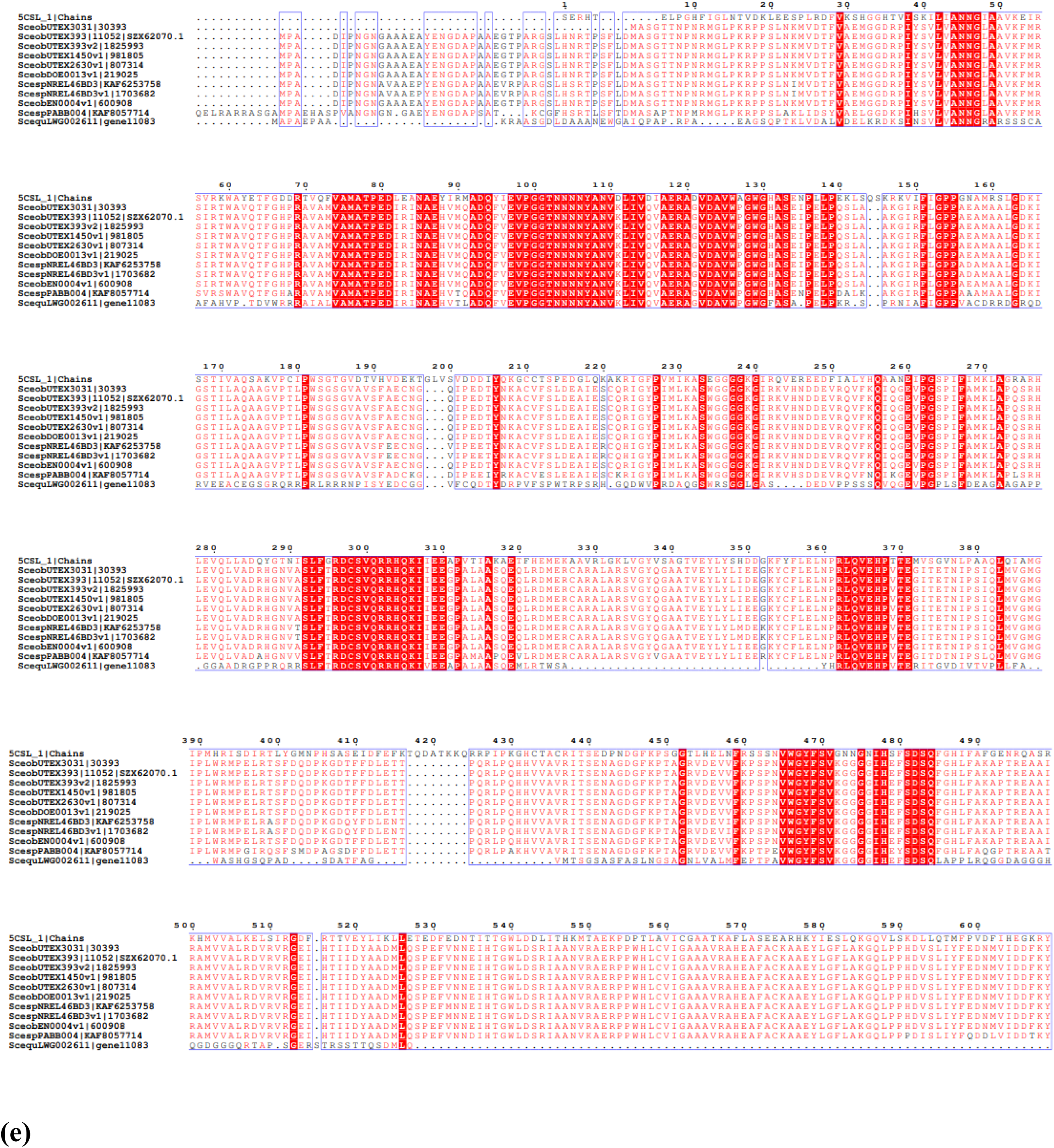
Multiple amino acid sequence alignments, performed using the Clustal Omega program and ESPript 3.0. The red colour represents the conserved regions. **(a)** Alignment between *Scenedesmus* BCs, 2VPQ (PDB entry) used as a template, red asterisk below the alignment indicates catalytic residues (K167, I208, K209, M219, H259), pink dots for glycine-rich loop, green up-triangle for the ERYV (Glu-Arg-Tyr-Val) motif; **(b)** Alignment between *Scenedesmus* BCCPs, 1BDO (PDB entry) used as a template, red asterisk above the alignment indicates catalytic residue, green dotted underline indicates the biotin-binding site, green asterisk for the biotin-binding residue (K186), pink up-triangle indicates AAAP (Ala-Ala-Ala-Pro) region. **(c)** Alignment between *Scenedesmus* α-CTs, 2F9I (PDB entry) used as a template; **(d)** Alignment between Scenedesmus β-CTs, 2F9I (PDB entry) used as a template, red asterisk below the alignment indicates catalytic Cysteine residues (C92, C95, C111, C114), other active site residues (S262, G270, G271, A248, G247, F293) are indicated by green asterisk. **(e)** Alignment between *Scenedesmus* homomeric ACCs, 5CSL (PDB entry) used as the template. The alignment was trimmed to show the protein identified for *S. quadricauda*.

**Fig 3:**
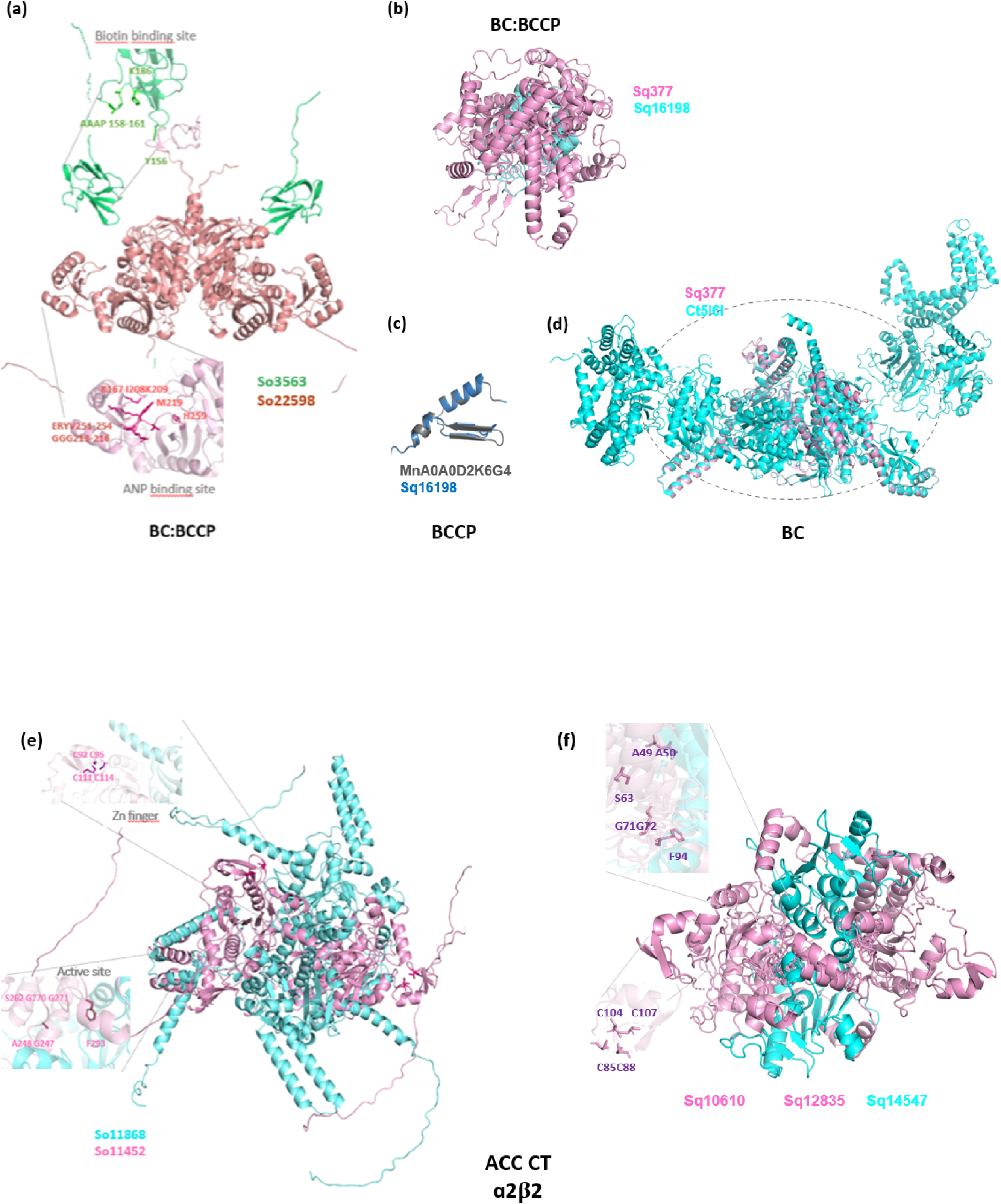
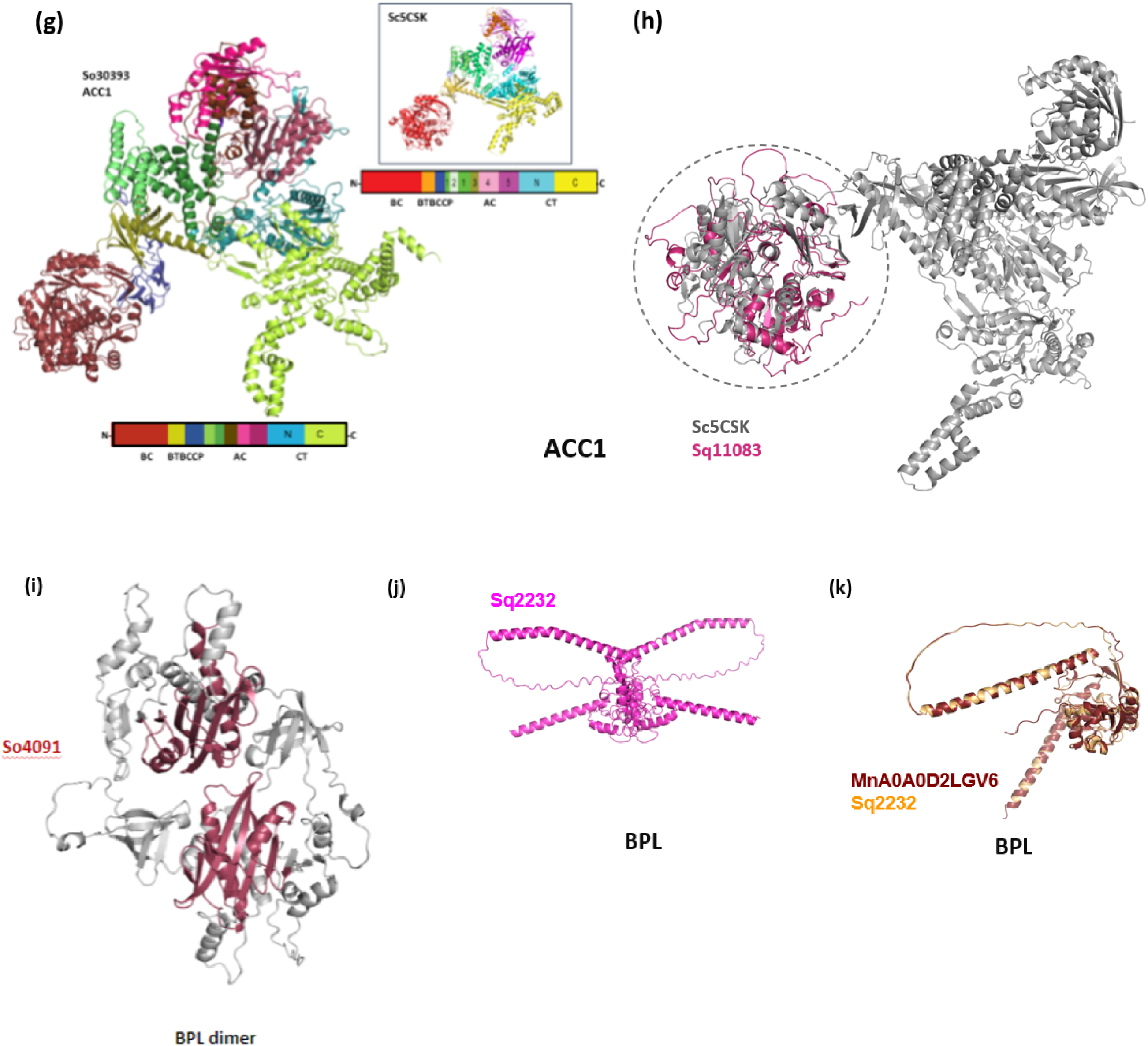
**(a)** Proposed model of the heterotetrameric BC:BCCP structure of ACC of S. obliquus UTEX B 3031. 3D modeling of Biotin Carboxyl Carrier Protein (green) and Biotin Carboxylase (dark pink). The biotin-binding amino acids are shown for one of the BCCPs in darker green **(b**) Proposed AlphaFold model of the heterodimeric BC:BCCP structure of *S. quadricauda* LWG002611. BCCP Sq16198 is shown in cyan and BC Sq377 in pink. **(c)** 3D model of BCCP Sq16198 (blue) and its superposition with *M. neglectum* A0A0D2K6G4 model (dark gray). **(d)** 3D model of BC Sq377 (pink) and its superposition with *C. thermophilum* 5I6I model (cyan). **(e)** Proposed model of the heterotetrameric α2β2 structures of ACC CT of S. obliquus UTEX B 3031 and **(f)** heterotetrameric α2β2 structure of ACC CT of *S. quadricauda* LWG002611. 3D modeling of Carboxyl Transferase α subunit (AccA) (cyan) and Carboxyl Transferase β Subunit (AccD) (pink) obtained with AlphaFold. Putative catalytic residues and atypic Cys4 ribbon Zn binding motif (zoom) are shown**. (g)** Proposed model of the cytosolic homomeric Acetyl CoA Carboxylase ACC1 of *S. obliquus* UTEX B 3031. 3D modeling shows from N to C-terminal: The BC domain in dark red, the BT domain in olive, the BCCP domain in dark blue, the Acc _central superfamily domains in green blue, brown, light and dark pink, and the carboxyl transferase CT domains in light blue and lime. The inset shows the crystal structure of the *S. cerevisiae* acetyl-CoA carboxylase holoenzyme 5CSK.**(h)** Proposed model of the cytosolic homomeric Acetyl CoA Carboxylase ACC1 of *S. quadricauda* LGW00611|11083 obtained with SWISS-MODEL. The inset shows the region of superposition between Sq11083 (dark Pink) and the crystal structure of the *S. cerevisiae* acetyl-CoA carboxylase holoenzyme 5CSK (gray)**. (i)** Proposed model of BPL dimer of S. obliquus UTEX3031 obtained with AlphaFold, in dark pink **(j)** Proposed 3D model of BPL of LWG002611 obtained with AlphaFold. **(k)** 3D SWISS-MODEL of BPL Sq2232 (orange) and its superposition with *M. neglect*um A0A0D2K6G4 model (burgundy).

The BCCP sequence of *S. quadricauda* (Sq16198), identified through genome analysis, was notably short and exhibited an inconclusive homology assignment with Phyre2. However, it showed homology with other BCCP from different algae inferred by orthoDB^27^ (https://www.orthodb.org/?gene=A0A0D2K6G4) and when a search for a template on SWISS-MODEL (http://swissmodel.expasy.org) resulted in a GMQE of 0.48% with the BCCP of another microalga *Monoraphidium neglectum* (A0A0D2K6G4.1.A) with a 72% of cover from residues 30-105 (Sq16198 numbering) (Fig. 3c). It’s noteworthy that the residues identified in the model BCCP structure from *Pedobacter heparinus* (1DBO, PDB code) as catalytic do not seem to be present in SqBCCP predicted sequence (Fig. 2b). In other *Scenedesmus* BCCP we have identified biotinyl/Lipoyl domain and biotin binding site (Fig. 1b, 2b).

Although BC and BCCP models obtained by SWISS-MODEL presented good superposition with their respective templates, when the 3D protein model of *S. quadricauda* LWG002611 BC:BCCP (Sq377: Sq16198) dimer was constructed with AlphaFold (Fig 3b), the IDDT per position score obtained was lower than 0.6%, and the model could not be superposed with any PDB structure from database.

On the other hand, a good BC:BCCP 3D model of *S. obliquus* UTEX3031 (So22598: So3563) was constructed with AlphaFold (Fig. 3a). Each pair of BC subunits showed contact between 3,0Å with the C-terminal region of each chain. The BCCP subunits in prokaryotes bind at 1:1 ratio or 1:2 ratio, the best relation found for this model was at 1:1 ratio. The biotin-binding residues were conserved and are shown for one of the BCCPs in dark green. In the 3D protein model of BC:BCCP we predicted conserved amino acid residues Lys167, Ile208, Lys209, Met219, His259 coordinated to the ATP molecule through electrostatic and hydrogen bonding interactions in ATP binding site. Glycine-rich loop known to play an important role in the enzyme’s catalytic function by conferring flexibility to adopt different conformations^27^ was observed located near the active site of biotin carboxylase. ERYV (Glu-Arg-Tyr-Val) motif identified in the 3D model possibly plays a role in the binding and recognition of biotin, as well as for the catalytic activity of the enzyme (Fig. 2a). Further experimental study and structural analysis are required for the confirmation of specific residues involved in ATP binding. Biotin binding site ‘GQTVCIIEAM**<u>K</u>**LMNEIEA’ having 90% homology with *E. coli* (PDB entry: 1BDO) has been detected in BCCP (Fig 2b, 3a). The biotin molecule would be covalently linked through an amide bond with a conserved lysine (**<u>K</u>**186) residue in the binding as reported previously in *E. coli*^28^ (Fig. 2b). A conserved sequence motif ‘AAAP’ (Ala-Ala-Ala-Pro) region detected in the 3D model functionally denotes the recognition site for the attachment of biotin^29^ (Fig. 2b, 3a). Biotinylated BCCP acts as a substrate for the subsequent carboxylation reactions catalyzed by the BC enzyme^30^.

It is interesting that both BC and BCCP sequences of *S. quadricauda* have not shown any similarity with prokaryotes^31^. The signal sequence of Sq377 suggested that the identified enzyme can be nuclear-encoded proteins and imported into chloroplasts. Some previous evidences are also available for algal BC and carboxyltransferase domains showing strong similarity with animal and yeast ACCases^32^. The similarity of Sq377 with eukaryotes may have some significance regarding the higher amount of lipid production^1,10^ than other *Scenedesmus* species. SqCT-β (Sq10610, Sq12835) was also shown similarity with other microalga rather than prokaryotes. Overall, the enzymes of heteromeric ACC of *S. quadricauda* were shown homology with prokaryotes and to some extent with eukaryotes, whereas enzymes of *S. obliquus* were shown homology completely with prokaryotes.

*S. quadricauda* CT α (Sq14547) sequence showed strong homology with *E. coli* and others Gram-negative bacterium revealing its prokaryotic origin (Table 2), whereas CT-β (Sq10610, Sq12835) showed an SWISS-MODEL GMQE 0.64 % with *M. neglectum* β-CT (A0A0D2KHA0.1.A) The detection of a carboxyl transferase domain in both the C-terminal α-CT (Fig. 1c) and the N-terminal β-CT (Fig. 1d), was achieved by analyzing by catalytic domain search. Furthermore, the conservation of amino acid residues in the C-terminal region for α-CT and in the N-terminal region for β-CT, as well as the conservation of the catalytic residues defined for 2F9I, was demonstrated by aligning the sequences with the corresponding 2F9I chains (Fig. 2c, 2d). We identified the residues corresponding to the Zn-binding sites (C85C88 C104C107) and the catalytic residues in Sq10610 (S63G71G72 F94A49A50). This statement was also shown in the AlphaFold 3D model of 2ɑ-CT2β-CT (Fig 3f). Besides, a 3D protein model of the heterotetrameric α2β2 structure of CT (So11868 and So11452) of *S. obliquus* UTEX3031 was constructed with AlphaFold (Fig. 3e). As occurs in 2F9I *S. aureus* crystallized heterotetramer, each pair of CT subunits presents a single active site, in which α-CT binds carboxybiotin and β-CT binds acetyl-CoA^33^. The structurally similar α and β monomers (cyan and pink) possess an α/β spiral core composed of a long, twisted, and tapered six-stranded mixed with six β sheet. The core is flanked by helical regions that move away from the structure. These helical elements in CT-β are interrupted by a Zn binding motif that is an atypical Cys4 ‘zinc ribbon’ with two antiparallel β sheets and a short α helix surrounding the four Cys, characteristic for β-CT subunit in prokaryotes and plants. We have detected catalytic Cysteine residues (C92, C95, C111 and C114) other active site residues (S262, G270, G271, A248, G247, F293) in β-CT component (Fig. 2d, 3e).

**Table 2:**
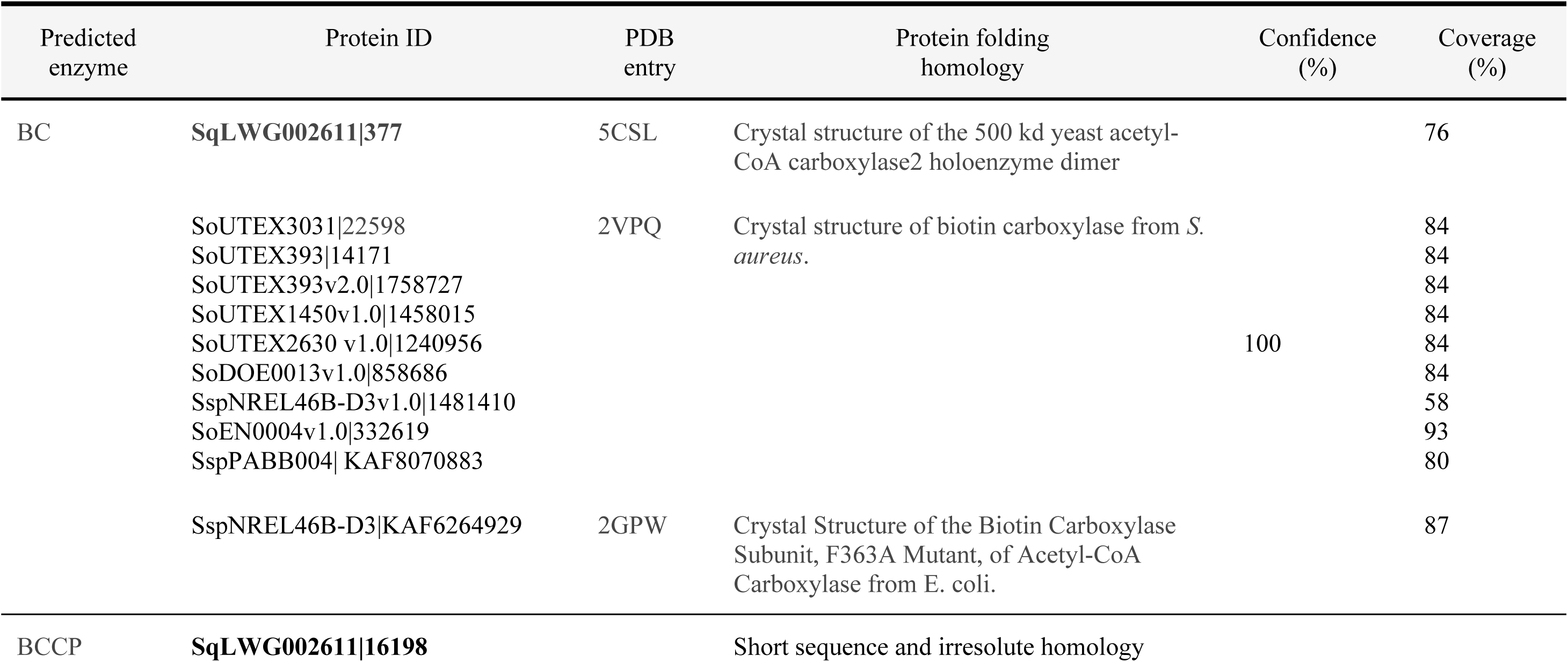
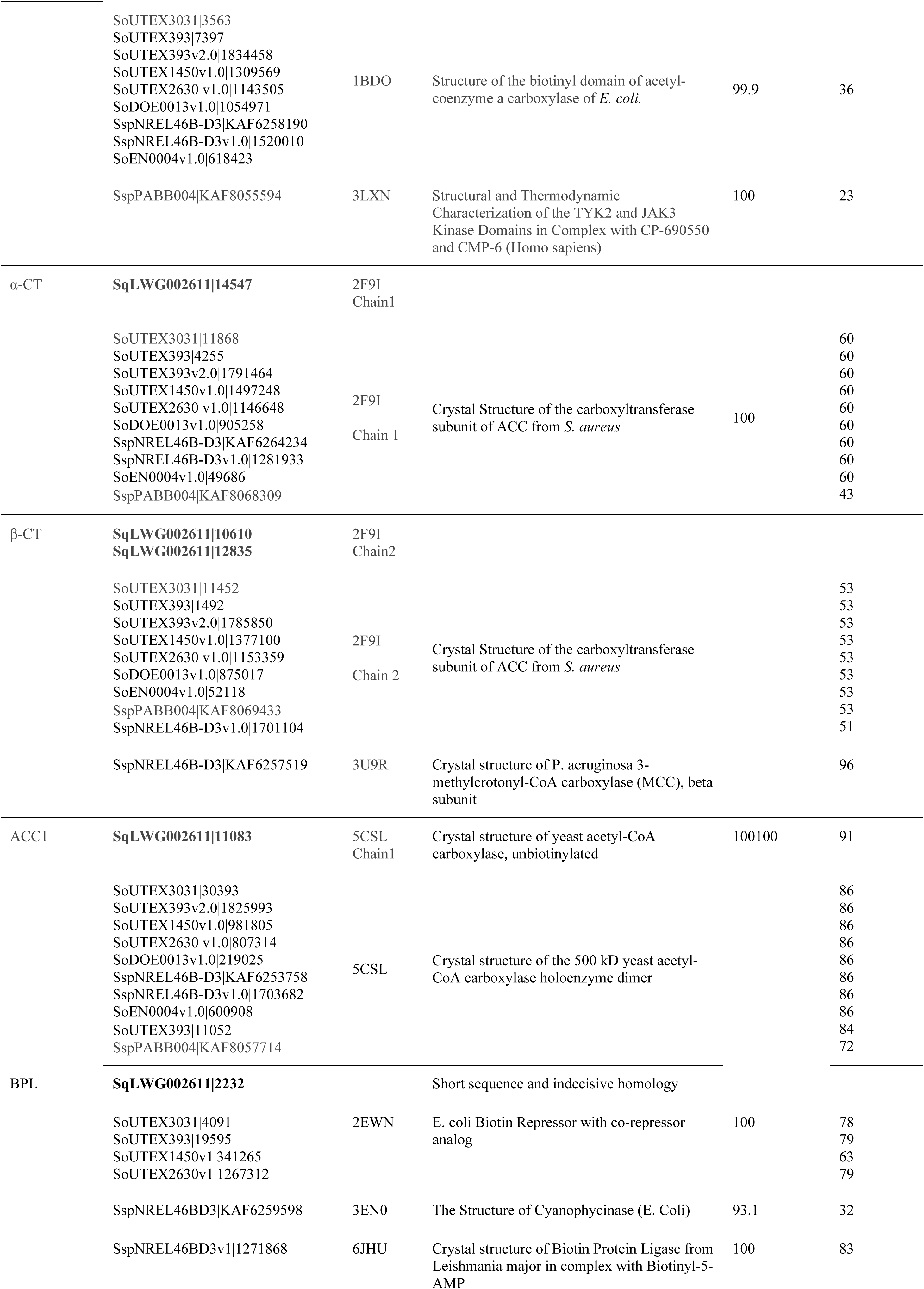
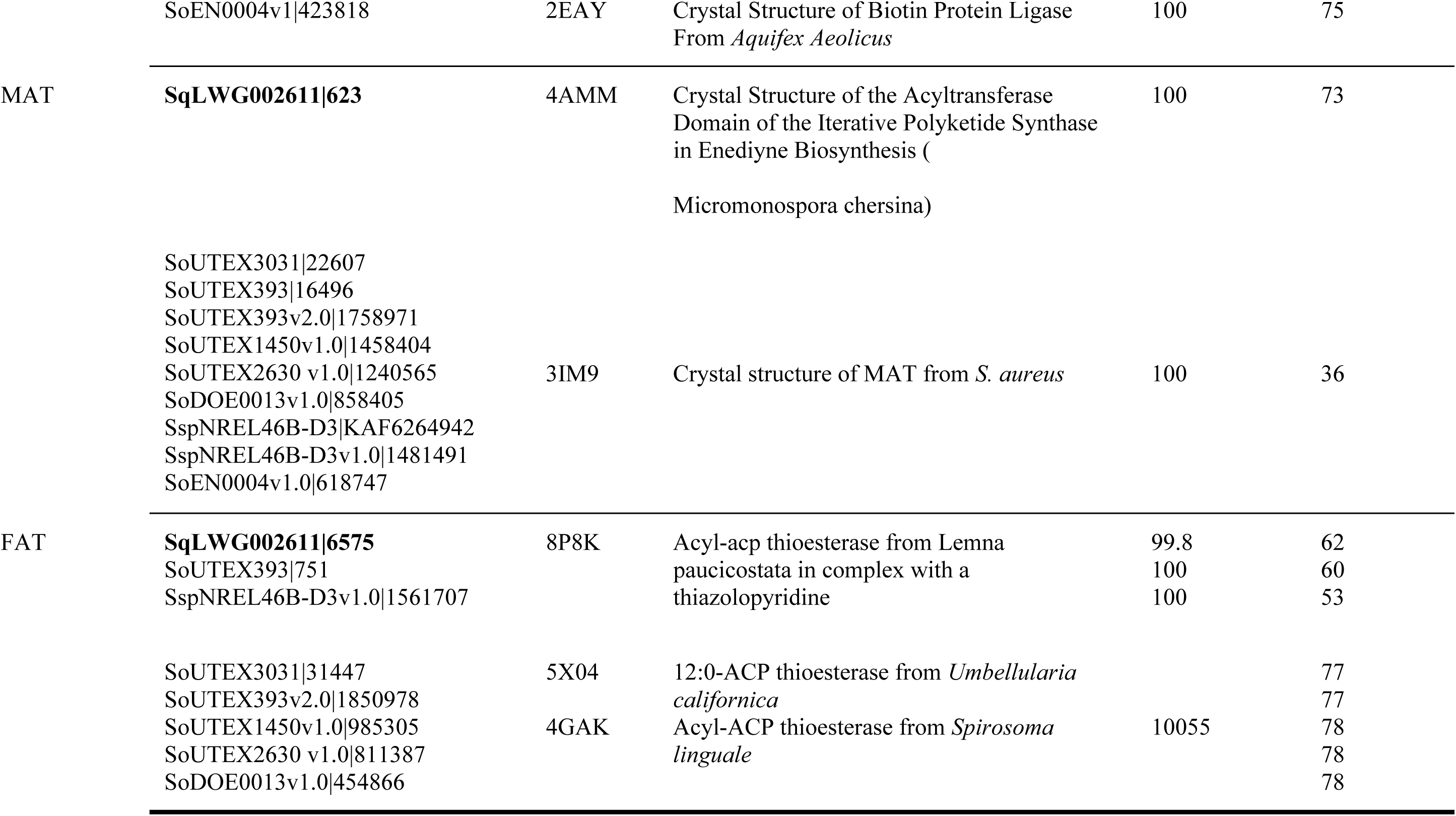
Comparative Phyre2 predicted protein folding homology of *S. quadricauda* LWG002611 and other *Scenedesmus* species.

Homomeric ACC of all the *Scenedesmus* analyzed, obtained a 100 % homology confidence score with eukaryotic yeast ACC1 reflecting its eukaryotic origin^29^. Only the BC domain seems to be present in the inferred sequence for *S. quadricauda* (Fig 1e and 3h), the possible existence of another alternative reading frame coding for the complete sequence does not escape our consideration, although the complete sequence needs to be studied experimentally. In homomeric ACC of *S. obliquus* UTEX 3031, all domains were found on a single peptide (Fig. 1e, 3g). Both N- and C-terminal biotin carboxylase domains interact with BCCP, and ATP grasp domain binds ATP which catalyzes the ATP-dependent carboxylation of the biotin cofactor, using bicarbonate as the CO_2_ donor^28^.

Absence of plastid signal peptide indicates that homomeric ACC is not located in chloroplast and it is probably located in cytosol as reported earlier^8^. This observation needs to be experimentally proven. The protein model of the homomeric So30393 ACC (*S. obliquus* UTEX3031) was also built with AlphaFold (Fig. 3g). Their domains showed a spatial arrangement similar to that of the crystallized structure of ACC from *S. cerevisiae* (PDB entry: 5CSL, 42.16% sequence identity and 80% cover)^34^. Biotin binding domains are positioned near the CT active site, and the BCCP domain is located near the center of the enzyme in both organisms (Fig. 3h). However, So30393 ACC presents some differences, being its BCCP domain longer than *S. cereviseae* and presents an additional ACC domain in its central region. Further studies are needed to determine the functional implications of these changes.

Despite the existence of the two ACC forms, published evidence and studies recognized the heteromeric ACCs present in algal plastids are majorly responsible for algal *de novo* FAs synthesis^8^. Whereas cytosolic ACCs are responsible for the elongation of FAs in the endoplasmic reticulum (ER)^6^ and the elongation of FAs to generate very long chain FAs as reported in *Arabidopsis*^7^.

We have also identified a biotin-protein ligase (BPL) enzyme in all *Scenedesmus* species that possibly mediates the covalent attachment of biotin to a specific lysine residue of BCCP and functions as a biotin operon repressor^21^ (Fig. 3i, 3j, 3k).

When the 3D protein model of *S. quadricauda* BPL (Sq2232) was constructed with AlphaFold as a dimmer (Fig 3j), the IDDT per position score was lower than 0.35% and the model could not be superposed with PDB structures from the database. Swiss-model searching for BPL identified *M. neglectum* BPL (A0A0D2LGV6.1.A) as the best template, with GMQE equal to 0.73 and 73.34 % identity and a coverage of 95 %. The superposition of these structures can be seen in Figure 3k and our sequence prediction matches the one indicated for that template. When the 3D protein model of the BPL structure (So4091) of *S. obliquus* UTEX3031 was constructed with AlphaFold, a dimeric structure resembling to *Staphylococcus aureus* (Sa)biotin protein ligase (PDB entry: 3RKX) was obtained (Fig. 3i). SaBPL is a dimeric class II BPL enzyme, with an additional DNA binding domain alongside its catalytic ligase domain. However, while SaBPL exhibits a side-by-side dimer structure, our SoBPL model is a domain-swapped dimer. Additionally, despite the presence of extra N and C-terminal domains in the model, a catalytic domain search conducted on NCBI did not assign any identity to them. Further experimental analysis is required to determine whether these domains belong to a new class of BPL structure.

### 3.2 Malonyl-CoA: acyl carrier protein transacylase (MAT)

A single Malonyl CoA: ACP transacylase (MAT) gene sequence was identified from reference-guided assembly of *S. quadricauda* LWG002611, showing homology with bacteria *Micromonospora chersina* (Table 2). We also analyzed the presence of MAT sequences in other selected *Scenedesmus* taken from PhycoCosm and NCBI showing homology with bacteria *S. aureus* (Table 1, 2). These results indicated that although all MAT showed bacterial homology, the *S. quadricauda* MAT enzyme differs from that of other *Scenedesmus* strains. This distinction was further explored through 3D modeling, aiming to identify differences in catalytic sites compared to other *Scenedesmus* strains. The acyl carrier protein transacylase domain was present in all sequences, whereas in Sq623, only the ACP binding domain was found (Fig 4a). Alignment between all *Scenedesmus* MAT sequences taken from PhycoCosm and NCBI revealed conserved motifs and catalytic sites (Fig. 5a).

**Fig 4:**
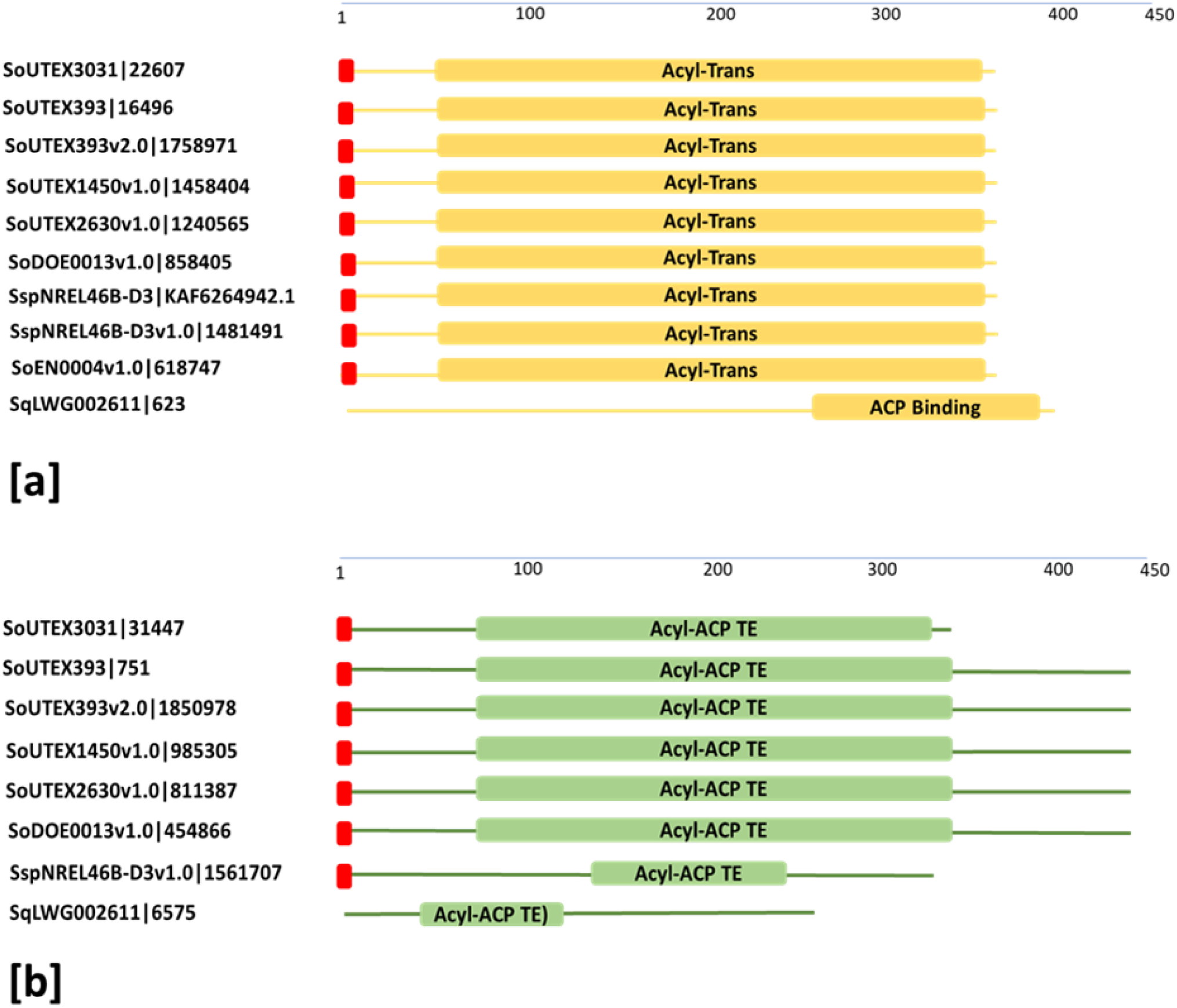
Protein domain architecture showing all predicted domains, red squares represent signal peptides. **(a)** Acyl transferase domains of Malonyl-CoA: acyl carrier protein transacylase (MAT) are represented by yellow rectangular bar; **(b)** Acyl ACP thioesterase domains of Fatty-acyl CoA Thioesterase (FAT) are represented by green rectangular bar.

**Fig 5:**
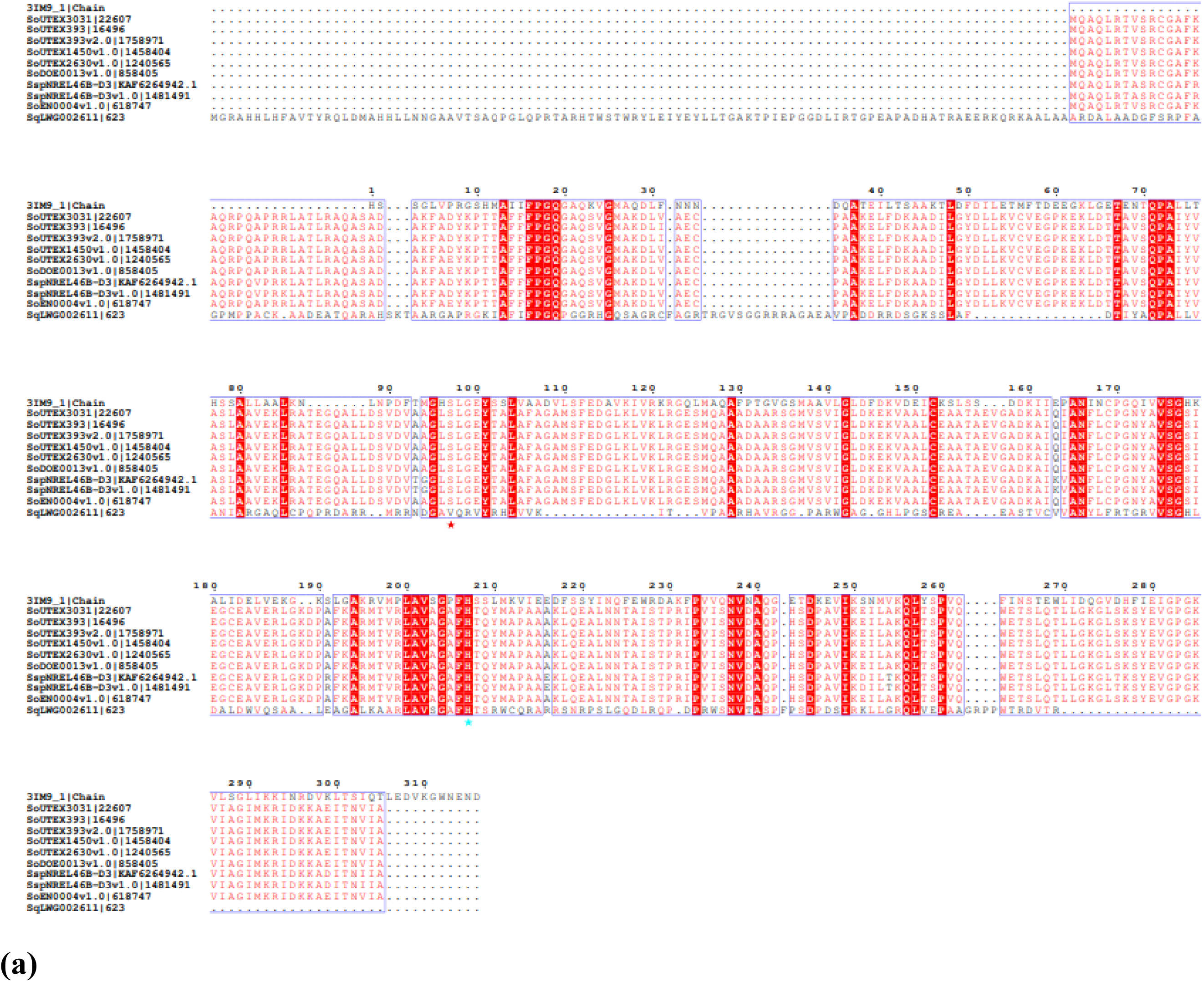
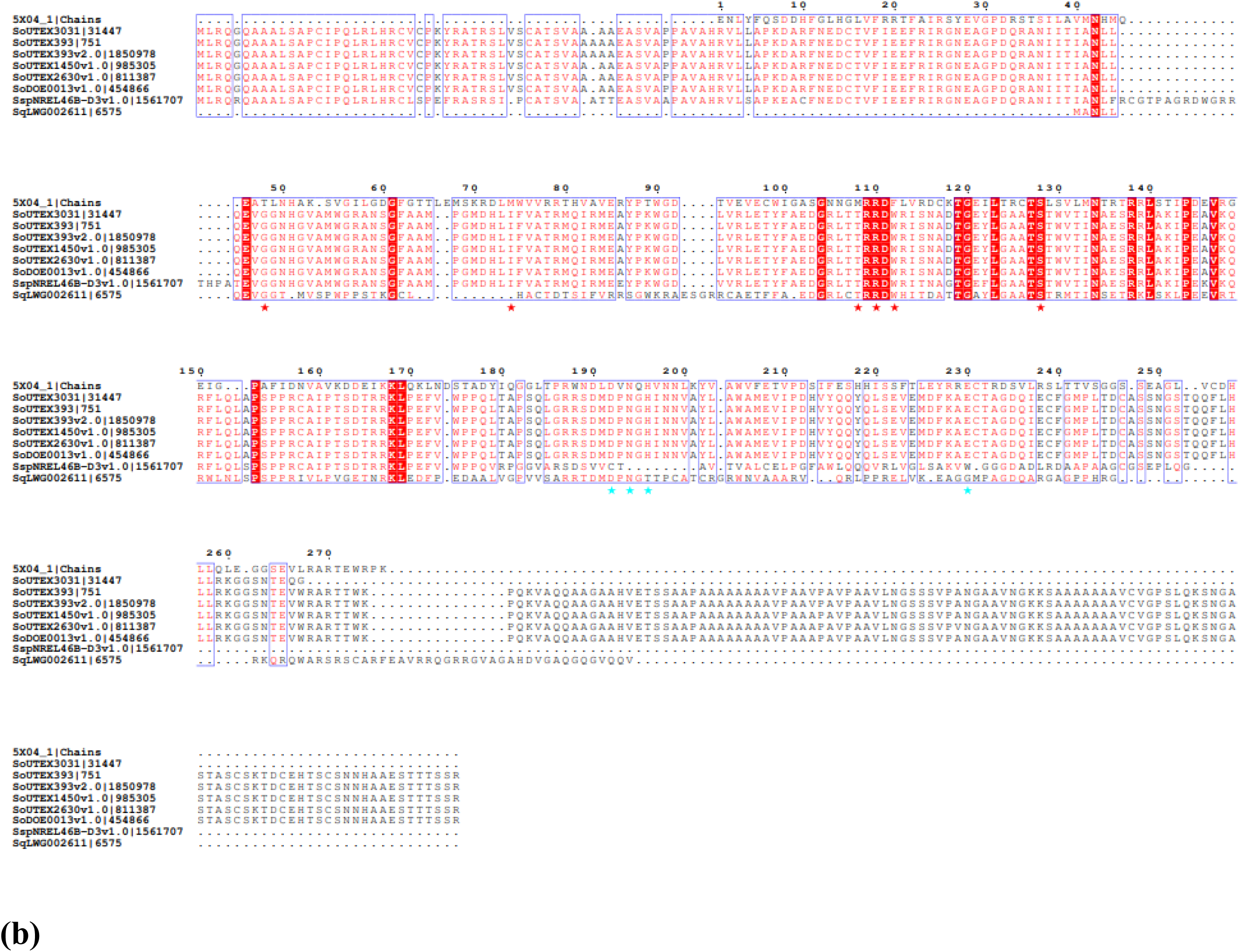
The multiple amino acid sequence alignment, performed using the Clustal Omega program and ESPript 3.0 showing the conserved sequences. **(a)** Alignment between *Scenedesmus* MATs, 3IM9 (PDB entry) used as a template, red asterisk below the alignment indicates catalytic serine residues (S137) and cyan asterisk indicates histidine residue (H252); **(b)** Alignment between *Scenedesmus* FATs, 5X04 (PDB entry) used as a template, red asterisk below the alignment indicates catalytic residues (G104, T129, T163, R165, W167, S183) and cyan asterisk indicates residues (D249, N251, H253, E287).

A 3D protein model of the MAT Sq623 was also constructed with Alpha Fold (Fig. 6b). This model exhibits two αβ subdomains as well. However, the large subdomain lacks two helices and two β-sheets at the C-termini. The superposition of the Sq623 model with MAT from *S. aureus* (PDB entry: 3IM9, 27.5% identity and 31% coverage) showed acceptable spatial conservation and presents the catalytic amino acid residue positioned in a similar spatial location to Ser97 and His 207 (*S. aureus* 3IM9 numbering).

**Fig 6:**
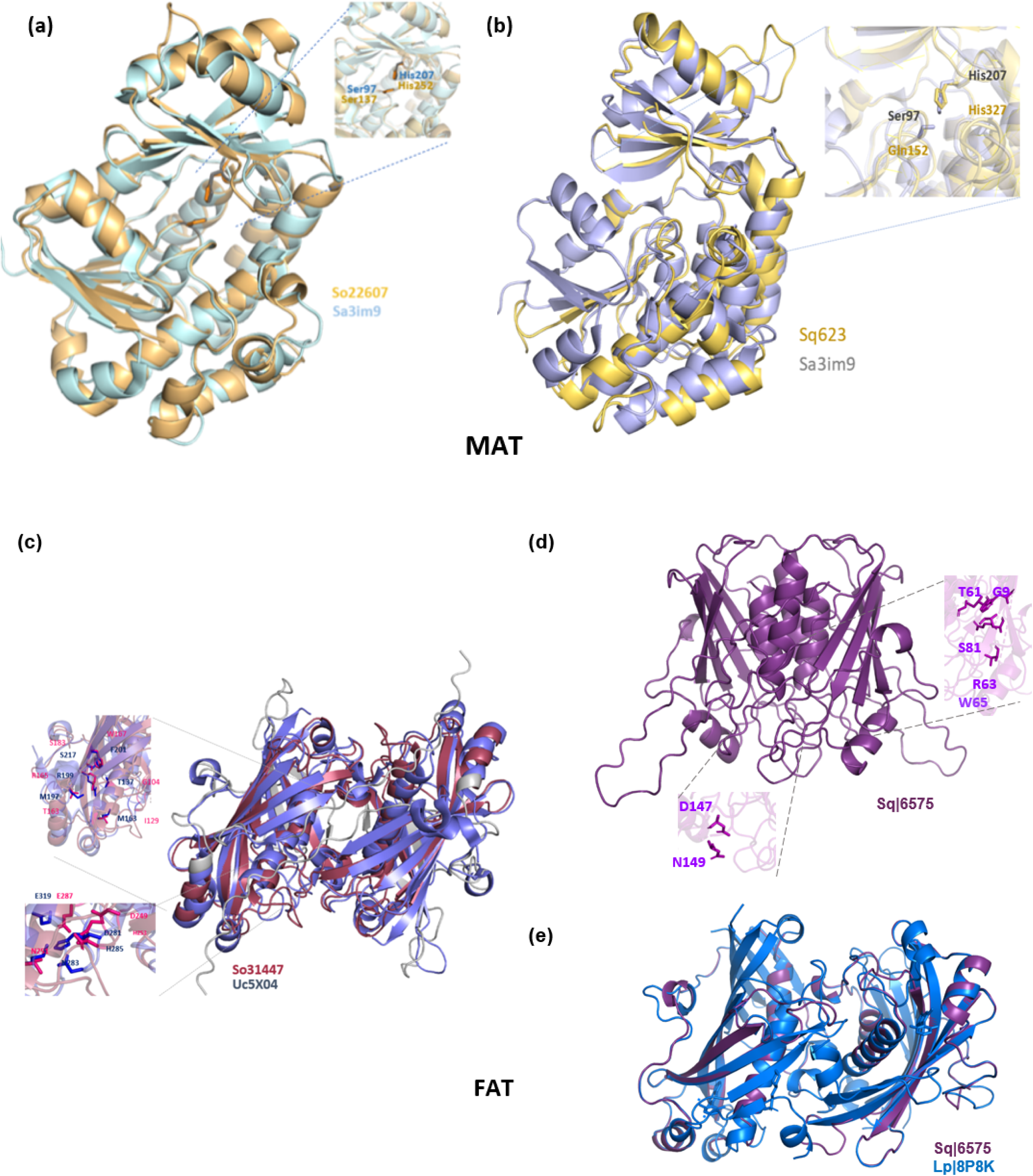
**(a)** AlphaFold 3D modeling of Malonyl CoA-acyl carrier protein transacylase for *S. obliquus* UTEX B 3031 (golden) and 3IM9 MAT from *S. aureus* (lightblue superposition). 3IM9 have two residues identified as active site which are conserved in Sceob3031|22607 MAT (zoom). (b) Superposition between AlphaFold 3D modeling of Sq623 (yellow) and the crystal structure of the MAT *S. aureus* enzyme 3IM9 (gray). **(c)** Proposed model of the dimeric structure of FAT *S. obliquus* UTEX B 3031. 3D modeling of AcylACP-thioesterase protein domain modeling for *S. obliquus* UTEX B 3031 (red) obtained with AlphaFold and 5X04 ACP thioesterase from Umbellularia californica (violet). Each pair of subunits have two hot-dog domains and a single active site; the dimeric structure is formed by the antiparallel bonding of the complete monomer structure (past to text). **(d)** Proposed 3D model of the dimeric structure of FAT of *S. quadricauda* (purple) obtained with AlphaFold. **(e)** Superposition between proposed 3D model of the dimeric structure of FAT of *S. quadricauda* Sq6575 (purple), obtained with SWISS-MODEL, and the crystal structure of the *L. paucicostata* 8P8K (blue).

3D protein model of the MAT So22607 was constructed with Alpha Fold (Fig. 6a). This structural model shows two αβ subdomains. The large one comprises a six β-stranded central β-sheet encircled by eleven helices while the small sub-domain comprises a four-stranded antiparallel β-sheet covered by two helices, similar to many characterized bacterial MAT enzymes. When the So22607 model was superposed with MAT from *S. aureus* (PDB entry: 3IM9, 36.27% identity and 86% coverage with So22607, they showed a good spatial conservation, and the catalytic amino acid residues Ser97 and His207 (*S. aureus* 3IM9 numbering) are positioned in a similar spatial location.

### 3.3 Fatty-acyl CoA Thioesterase (FAT)

One palmitoyl-acyl carrier protein thioesterase (FAT) gene sequence was identified in reference assembly *S. quadricauda* LWG002611 (Table 1, 2). Acyl ACP thioesterase domain was found in all selected sequences (Fig 4b). Interestingly FAT sequences had a very diverse structural homology. By analyzing with Phyre2 the maximum homology was found with sequences from plant *Lemna paucicostata* and *Umbellularia californica* followed by bacteria *Spirosoma linguale* (Table 2). By aligning all *Scenedesmus* FAT sequences taken from PhycoCosm and NCBI conserved motifs were determined (Fig. 5b).

Acyl-ACP thioesterase family/domain contains a duplication of two 4 hydroxybenzoyl-CoA thioesterase like domains (4HBT), which forms a homotetramer with four active sites. Each subunit of the 4HBT tetramer adopts a “hot-dog” fold. This domain contains a seven-stranded antiparallel β-sheet, which encloses a five-turn α-helical.

The Sq6575 3D model obtained with AlphaFold exhibits only two HBT domains and in a flipped orientation (fig 6d). On the other hand, the model obtained with SWISS-MODEL also exhibits only two HBT domains but presents the same orientation than So31447. When it was superposed with *Lemna paucicostata* FAT template (PDB entry: 8P8K, 32 % identity and 68% coverage with Sq6575), they showed partial spatial conservation and catalytic amino acids are conserved (Fig. 6d). However, residues E319 and H285 (Uc5X04 numbering) were not found to be conserved.

When the So31447 model was superposed with FAT from *Umbellularia californica* (PDB entry: 5X04_A, 30.91% identity and 80% coverage with So31447), they showed a good spatial conservation, and the catalytic and substrate binding amino acid residues (Fig. 6c) are positioned in a similar spatial location. This suggests that the assignment of the function of this protein is adequate. Catalytic residues were identified in the 3D model (G104, T129, T163, R165, W167, S183 D249, N251, H253, E287).

In summary, the high conservation of structure and amino acid sequence of the fatty acid synthesis enzymes from the analyzed *Scenedesmus* species suggests the preservation of this pathway in *Scenedesmaceae* family. It has been perceived that initially prokaryotes entered as endosymbionts and further incorporated there as an organelle by transferring genes to the genome of the host during evolution^31^. However, some contrasting results were also observed between *S. quadricauda* and other species of *Scenedesmus*. Maybe due to the unique sequences and binding sites of some enzymes, lipid production is higher in *S. quadricauda* than the other species as evident by our previous reports^1,10^. Although the knowledge of the *Scenedesmus* genome and transcriptome has increased in recent years, the biochemistry of its many enzymatic pathways has still been poorly characterized.

Understanding the catalytic sites and domains of fatty acid metabolism enzymes in microalgae is key to evaluating the potential of promising strains for biofuel production.

## Acknowledgments

This work is supported by the Agencia Nacional de Promoción Científica y Tecnológica (Grant No. ANPCyT PICT2019-00091); Consejo Nacional de Investigaciones Científicas y Técnicas (CONICET); Science and Engineering Research Board (SERB) (Grant No. SPG/2021/001796), New Delhi and Amity University Uttar Pradesh, Lucknow Campus; Authors are thankful to Msc Mariano Torres Manno for his help with 3D protein modelling; grateful to CSIR-National Botanical Research Institute, Lucknow for organism *S. quadricauda* LWG002611.

## Author information

HKS is a doctoral fellow from Amity University Uttar Pradesh.

MBV and NM are doctoral fellows from CONICET.

JB and MVB are research members from CONICET.

CND is assistant professor from Amity University Uttar Pradesh, Lucknow, India.

## Corresponding authors

Correspondence to Dr. Dasgupta and Dr. Barchiesi

## Author contributions

HKS, sequence searched, aligned and analyzed, MBV, protein modeling and analyses; NM, sequence analyses; MVB, project administration, wrote the manuscript; JB; analyzed the data, conceptualization, wrote manuscript; CND, conceptualization, interpreted the data, wrote the manuscript

